# Human TRPA1 is an inherently mechanosensitive bilayer-gated ion channel

**DOI:** 10.1101/2020.03.05.979252

**Authors:** Lavanya Moparthi, Peter M. Zygmunt

## Abstract

The Transient Receptor Potential Ankyrin 1 (TRPA1) channel is an intrinsic chemo- and thermo-sensitive ion channel with distinct sensory signaling properties. Although a role of TRPA1 in mammalian mechanosensory transduction *in vivo* seems likely, it remains to be shown that TRPA1 has the inherent capability to respond to mechanical stimuli. Here we have used the patch-clamp technique to study the response of human purified TRPA1 (hTRPA1), reconstituted into artificial lipid bilayers, to changes in bilayer pressure. We report that hTRPA1 responded with increased single-channel open probability (P_o_) within the applied pressure interval of 7.5 to 60 mmHg with a half maximum P_o_ (P_50_) value of 38.0 ± 2.3 mmHg. The P_o_ value reached a maximum close to 1 (0.87 ± 0.02) at 60 mmHg. Within the same pressure interval, hTRPA1 without its N-terminal ankyrin repeat domain (Δ1-688 hTRPA1) responded fully opened (0.99 ± 0.01) at 60 mmHg and with a P_50_ value of 39.0 ± 1.1 mmHg. The pressure-evoked responses of hTRPA1 and Δ1-688 hTRPA1 at 45 mmHg were inhibited by the TRPA1 antagonist HC030031, and the activity of purified hTRPA1 at 45 mmHg was abolished by the thiol reducing agent tris(2-carboxyethyl)phosphine (TCEP). In conclusion, hTRPA1 is an inherent mechanosensitive ion channel gated by force-from-lipids. The hTRPA1 mechanosensitivity is dependent on the redox environment, and it is suggested that oxidative stress shifts hTRPA1 into a protein conformation sensitive to mechanical stimuli.

## Introduction

The discovery of transient receptor potential (TRP) channels involved in mammalian sensory signaling has introduced a new class of polymodal proteins as detectors of chemical, temperature and possibly mechanical stimuli (1–4).

The finding that mammalian TRPA1 is activated by more than hundred different chemical compounds including thiol-reactive electrophilic compounds and oxidants as well as non-electrophilic compounds such as menthol and cannabinoids has consolidated TRPA1 as a unique chemosensor (5, 6). The original discovery of TRPA1 as a noxious cold sensor has been much debated but gained a lot of support over time (5, 6). Furthermore, TRPA1 has recently been revealed as an intrinsic warmth/heat receptor that may contribute to sensing warmth and uncomfortable heat in mammalians (5).

The proposal by David Corey and colleagues that mouse TRPA1, with its large intracellular N-terminal ankyrin repeat domain (N-ARD), could be a mechanosensor involved in hearing has triggered a great interest in TRPA1 as a potential primary mechanosensor within the mammalian sensory nervous system (4–6). Although a role for TRPA1 in mammalian normal and nociceptive signaling in response to mechanical stimuli seems convincing, no evidence of TRPA1 intrinsic mechanosensitivity has been provided (3, 5–7).

In this study, we have taken advantage of our experience from studies on the inherent chemo-and thermo-sensing properties of purified hTRPA1 when reconstituted into artificial lipid bilayers (8–10), to explore the mechanosensitivity of purified hTRPA1. In our previous studies, we noticed the impact of the redox environment on the ability of TRPA1 to respond to temperature (9), which is perhaps not surprising as mammalian TRPA1 has many cysteine residues (e.g., 28 in hTRPA1 and 31 in mouse TRPA1) that are involved in molecular intra-and inter subunit disulfide interactions as well as being targeted by thiol modifying agents (5, 6). We therefore also investigated the effect of the thiol reducing agent tris(2-carboxyethyl)phosphine (TCEP) on hTRPA1 mechanical activity.

## Results

The hTRPA1 with and without its N-terminal ARD (Δ1-688 hTRPA1) were purified and reconstituted into artificial lipid bilayers for electrophysiological recordings of single-channel activity in response to various negative pressures in a stepwise manner (step width 10 mbar). When exposed to an increase in pressure, both hTRPA1 (Fig. 1) and Δ1-688 hTRPA1 (Fig. 2) responded with increased single-channel activity. Within the applied pressure interval of 7.5 to 60 mmHg, the single-channel open probability (P_o_) value reached a maximum close to 1 (0.87 ± 0.02) at 60 mmHg with a half P_o_ (P_50_) value of 38.0 ± 2.3 mmHg for hTRPA1 (Fig. 1A). Δ1-688 hTRPA1 were fully opened (0.99 ± 0.01) at 60 mmHg with a P_50_ value of 39.0 ± 1.1 mmHg (Fig. 2A). The TRPA1 antagonist HC030031 blocked both hTRPA1 and Δ1-688 hTRPA1 single-channel activity at 45 mmHg (Fig. 1B and Fig. 2B). This concentration of HC030031 also inhibited hTRPA1 and Δ1-688 hTRPA1 single-channel activity evoked by chemical ligands and cold/warm temperatures (9, 10). The single-channel activity of hTRPA1 at 45 mmHg was abolished by the thiol reducing agent TCEP (Fig. 1C) at a concentration that abolished hTRPA1 single-channel activity evoked by H_2_O_2_ and cold/warm temperatures (9). No channel activity was observed in membranes without the purified hTRPA1 proteins within the applied pressure interval of 7.5 to 60 mmHg (not shown).

**Figure 1.**
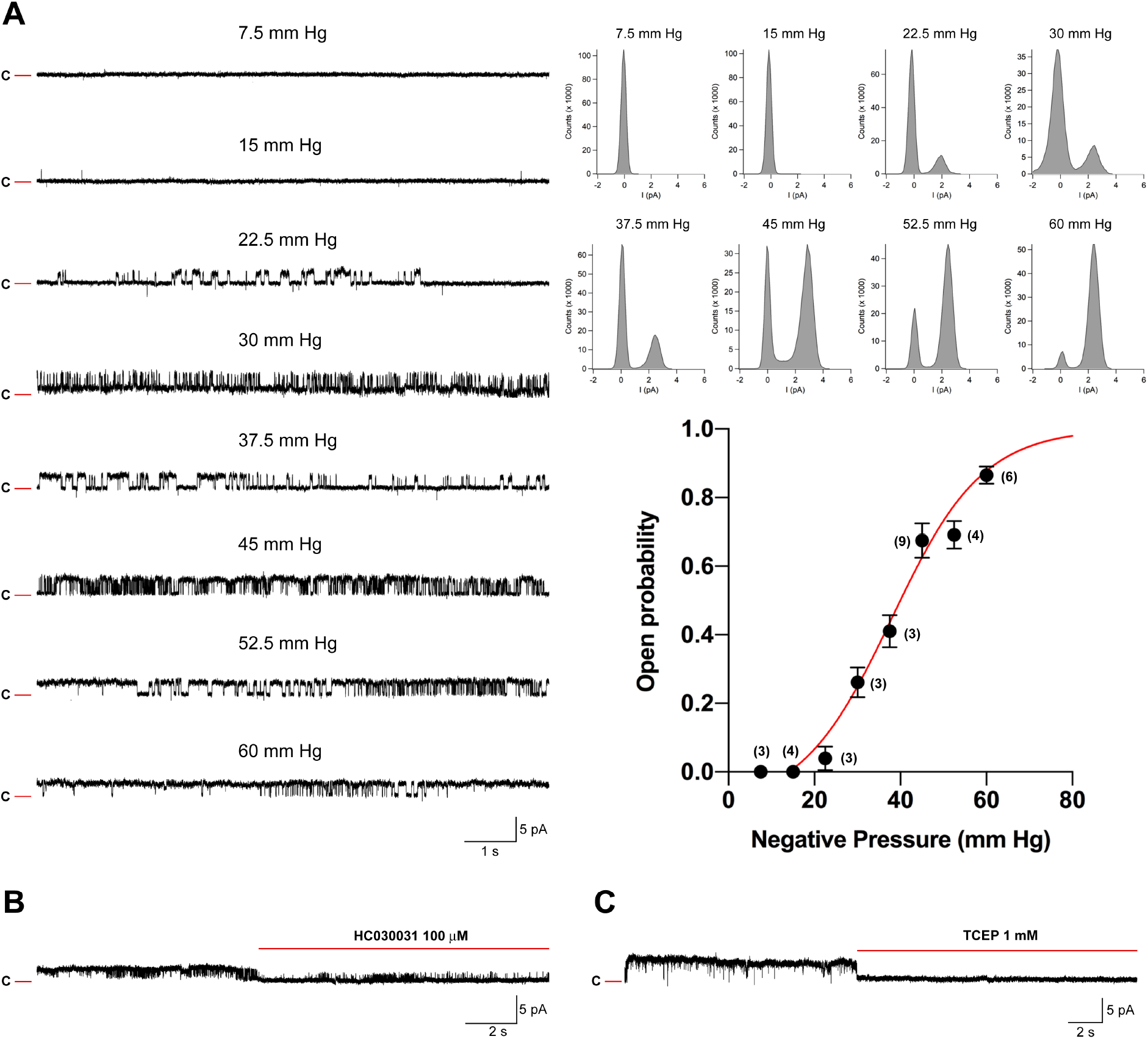
Human TRPA1 is intrinsically mechanosensitive. Purified hTRPA1 was reconstituted into planar lipid bilayers and single-channel currents were recorded with the patch-clamp technique in a symmetrical K^+^ solution. (A) As shown by representative traces and corresponding amplitude histograms, exposure to a change in negative pressure evoked outward single-channel currents at a test potential of +60 mV. The graph shows single-channel mean open probability values as a function of applied negative pressure. Data were fitted with a Boltzmann equation and each data point is the mean ± SEM of 3-9 observations (shown within parentheses). (B) Trace showing inhibition of hTRPA1 mechanical activity at 45 mmHg by the TRPA1 antagonist HC030031 (n=2). (C) Trace showing inhibition of hTRPA1 mechanical activity at 45 mmHg by the thiol reducing agent TCEP (n=3). C = closed-channel state, upward deflection = open channel state.

**Figure 2.**
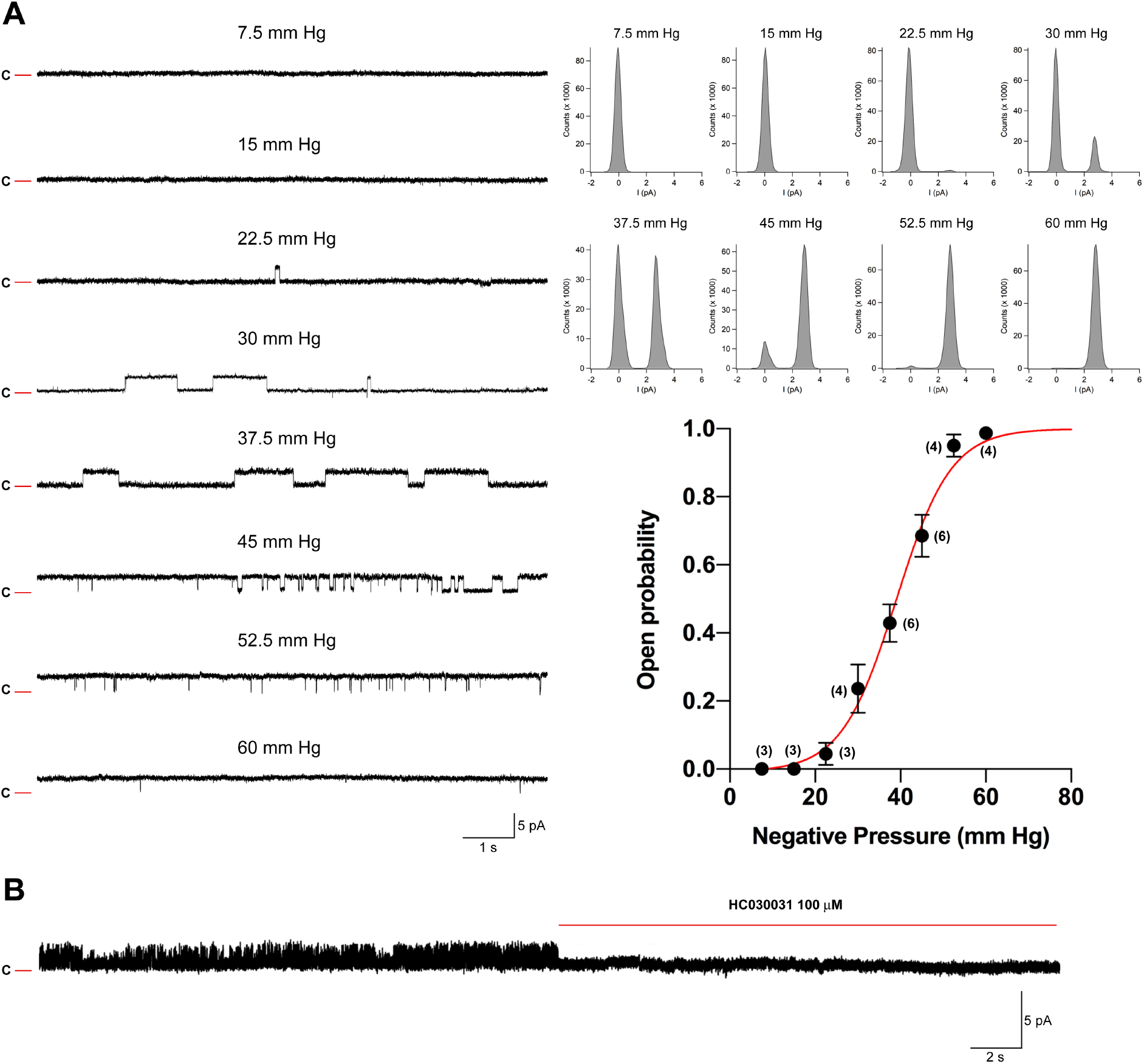
Human TRPA1 without its N-terminal ankyrin repeat domain is intrinsically mechanosensitive. Purified Δ1-688 hTRPA1 was reconstituted into planar lipid bilayers and single-channel currents were recorded with the patch-clamp technique in a symmetrical K^+^ solution. (A) As shown by representative traces and corresponding amplitude histograms, exposure to a change in negative pressure evoked outward single-channel currents at a test potential of +60 mV. The graph shows single-channel mean open probability values as a function of applied negative pressure. Data were fitted with a Boltzmann equation and each data point is the mean ± SEM of 3-6 observations (shown within parentheses). (B) Trace showing inhibition of hTRPA1 mechanical activity at 45 mmHg by the TRPA1 antagonist HC030031 (n=2). C = closed-channel state, upward deflection = open channel state.

In the same automated planar patch-clamp system, reconstitution of the purified bacterial mechanosensitive channel of large conductance (MscL) and the purified mammalian two-pore domain potassium channel TREK-2 displayed graded single-channel activity in response to changes in negative pressure within 60 - 100 mmHg for MscL and 30 - 90 mmHg for TREK-2 (11, 12).

## Discussion

A number of TRP channels has been suggested to be involved in mammalian mechanosensation under normal physiological conditions as well as in pathophysiology (2, 4–6). Among the suggested TRP channels, there is substantial evidence of TRPA1 being involved in mechanical sensory stimulation, and especially in noxious mechanotransduction e.g., related to nerve-injury, inflammation and anti-cancer treatment (5, 6). However, it still remains to be demonstrated that TRPA1 exhibit mechanosensitivity in the absence of extra-and intra cellular components (3, 5, 13).

In this study, we explored the possibility that hTRPA1 is an inherent mechanosensitive ion channel when purified and reconstituted into artificial lipid bilayers, using the same experimental conditions as in our studies consolidating hTRPA1 as an inherent chemo-and thermo-sensitive ion channel (8–10). We found that hTRPA1 without its N-ARD (Δ1-688 hTRPA1) displayed a similar mechanosensitive profile as intact hTRPA1, and thus the many N-terminal ankyrin repeats are not needed for TRPA1 to respond to mechanical stimuli as generally believed (4–6). However, this does not exclude a role of the N-ARD in coordinating mechanical stimuli and channel activity in a native environment by interacting with lipids of the cell membrane, as well as tethered with the cytoskeleton or the extracellular matrix. The association of N-ARD with proteins including TRPV1, fibroblast growth factor receptor 2 and prolyl hydroxylases (5, 6) may also influence TRPA1 mechanosensory signaling properties. Furthermore, any interaction with N-ARD cysteines by electrophilic compounds and redox agents could easily change the overall protein conformation (5, 6, 14) possibly affecting the mechanosensitivity of TRPA1. Indeed, here we found that the mechanical activity of hTRPA1, which is partially oxidized (9), was abolished by the thiol reducing agent TCEP that also neutralized the U-shaped thermosensitivity of purified hTRPA1 (9). Oxidants including H_2_O_2_ also sensitize TRPA1 heat responses in cells and isolated tissues (9, 15), and a recent study showed H_2_O_2_ sensitization of TRPA1-mediated mechanical responses in mouse trigeminal sensory neurons when exposed to a hyperosmotic solution (16). Taken together, it is suggested that oxidative stress and perhaps other environmental factors (e.g., pH, Ca^2+^, polyphosphates) alone or in combination can shape the response to mechanical stimuli by shifting hTRPA1 into a force-to-lipid sensitive protein conformation. Thus, the concerns raised regarding experimental condition and TRPA1 thermosensitivity (5, 6) may also be valid in studies of TRPA1 mechanosensitivity, and explain why TRP channel mechanical activity cannot always be detected at a cellular level or in artificial lipid bilayers (7).

Another interesting aspect on TRPA1 mechanosensation is that the effect of non-electrophilic TRPA1 ligands may be indirect by changing the lipid tension stress on TRPA1 within the cell membrane (17). For example, the ability of primary and secondary alcohols as well as different alkyl-substituted phenol derivatives including carvacrol to activate hTRPA1 increased with increasing lipophilicity (17). Our observation that carvacrol induced conformational changes of purified hTRPA1 in a bilayer-free environment indicates, however, a direct interaction with TRPA1 (9). In our study on the structure-activity relationship of cannabinoids and TRPA1 from mouse and human, methylation of the Δ^9^-tetrahydrocannabinol C-1 hydroxyl group removed its ability to activate TRPA1 although both compounds had very similar lipophilicity (18). Furthermore, the synthetic cannabinoid and very potent cannabinoid CB1/CB2 receptor agonist CP55940 (logP = 6.2) did not activate purified hTRPA1 in bilayer recordings whereas the subsequent exposure to the less lipophilic Δ^9^-tetrahydrocannabinol (logP = 5.5) activated hTRPA1 (10). The modifications of the Δ^9^-tetrahydrocannabinol alicyclic C-9 methyl group and the C-3 carbon tail further suggested a specific cannabinoid binding site on TRPA1 (18). Nevertheless, future studies of the lipid-TRPA1 interaction will be of great interest, as a change in membrane fluidity could still be part of the pharmacology of non-electrophilic TRPA1 ligands including cannabinoids.

In summary, hTRPA1 is an inherent mechanosensitive channel that like the other mammalian mechanosensitive channels TREK-1, TREK-2, TRAAK and Piezo 1 is gated by force-from-lipids. The hTRPA1 mechanosensitivity is dependent on the redox environment, and it is suggested that oxidative stress shifts hTRPA1 into a protein conformation sensitive to mechanical stimuli. Future studies of TRPA1 from both vertebrates and invertebrates could help to obtain an evolutionary detailed mechanistic understanding of the intrinsic mechanosensitive properties of TRPA1 and perhaps other TRP channels.

## Materials and Methods

The expression and purification of hTRPA1 were performed as described previously (10). Purified hTRPA1 was reconstituted into preformed planar lipid bilayers composed of 1,2-diphytanoyl-sn-glycero-3-phosphocholine (Avanti Polar Lipids) and cholesterol (Sigma-Aldrich) in a 9:1 ratio and produced by using the Vesicle Prep Pro Station (Nanion Technologies). Under these conditions, a uniform protein orientation is favored with N- and C-termini facing the recording chamber (i.e., the “cytosolic compartment”) (10). Ion channel activity was recorded using the Port-a-Patch (Nanion Technologies) at a positive test potential of +60 mV in a symmetrical K^+^ solution (50 mM KCl, 10 mM NaCl, 60 mM KF, 20 mM EGTA, and 10 mM Hepes; adjusted to pH 7.2 with KOH) and at room temperature (20°-22°C). Negative pressure was applied in a stepwise manner (step width 10 mbar) with the suction control pro (Nanion Technologies) to evoke TRPA1 currents. Signals were acquired with an EPC 10 amplifier and PatchMaster software (HEKA) at a sampling rate of 50 kHz. Electrophysiological data were analyzed using Clampfit 9 (Molecular Devices) and Igor Pro (WaveMetrics). Data were processed by a Gaussian low-pass filter at 1000 for analysis and 500 Hz for traces. The single-channel mean open probability (P_o_) was calculated from time constant values, which were obtained from exponential standard fits of dwell time histograms.

## Acknowledgements

This study was supported by the Swedish Research Council (2014-3801) and the Medical Faculty of Lund University – ALF (Dnr. ALFSKANE-451751).

